# The small GTPase Rem2 modulates sex-dependent spatial learning by regulating CA1 glutamate receptor composition

**DOI:** 10.64898/2026.09.15.751881

**Authors:** William E. Brew, Hassan E. Mohammed, Afrina Asad Meghla, Serendipity Coniglio, Eleanor Labriola, Regan Skudlarek, Kishore Kumar S. Narasimhan, Shashank M. Dravid, Gillian Queisser, Anna R. Moore, Victor M. Luna

## Abstract

The small GTPase Rem2 is a key signaling molecule involved in synaptic formation, dendritic complexity, spine formation, and regulation of CaMKII-dependent long-term potentiation **(LTP)**. However, it remains unclear how Rem2 explicitly impacts learning and memory. To address this issue, we deleted Rem2 specifically in dorsal CA1 neurons of male and female mice and assessed spatial learning using a spatial object recognition **(SOR)** task. We found that males outperform females in this behavioral assay and that deleting Rem2 did not impact spatial learning in males. In contrast, Rem2 deletion significantly improved spatial learning in females enabling them to perform as well as males in the SOR task. Using an automated Western blot system to assess all known AMPA- and NMDA-mediated glutamate receptor (AMPAR and NMDAR) subunits in each mouse, we found that the sex-dependent change in SOR we observed was likely due to Rem2 increasing GluN2D expression in the interneurons of female mice only. Furthermore, using S-statistics to compare the overall AMPAR and NMDAR composition among groups, we found that males and females normally have divergent covariance structures but this divergence is eliminated when Rem2 is deleted from CA1 neurons. These results reveal an unexpected role of Rem2 in maintaining sex differences in glutamate receptor composition and spatial learning abilities. To our knowledge, this is the first demonstration of a signaling molecule that confers sexual dimorphism to excitatory synapses. As such, Rem2 may play a critical role in understanding how sex-dependent symptoms of neurodevelopmental and neurodegenerative disorders arise.

## 1 INTRODUCTION

Synapses are the neurobiological bases for all cognitive functions. Determining the signaling molecules that regulate synaptic formation are thus critical to our basic understanding of learning and memory as well as to the molecular events that lead to their dysfunction.

Rem2, a small GTPase from the RGK subfamily of Ras-like GTPases, was previously identified as a key component of excitatory synapse formation via activity-dependent RNA screening (Paradis et al., 2007). Rem2 has since been implicated in several processes critical to synaptic plasticity including dendritic branching, spine formation, and synapse development (Ghiretti and Paradis, 2011; Ghiretti and Paradis, 2014; Ghiretti et al., 2014; Moore et al., 2013). Rem2 has also been found to interact with CaMKII as both a phosphorylation substrate and an endogenous inhibitor of CaMKII (Ghiretti et al., 2013; Royer et al., 2018) implicating Rem2 in calcium-dependent signaling pathways critical to synaptic plasticity. In addition, Rem2 has been shown to modulate gene expression in response to neuronal depolarization (Kenny et al., 2017) further supporting its role as an activity-regulated transcriptional modulator. Recently, Rem2 has been revealed as a negative regulator of long-term potentiation **(LTP)** in the hippocampus acting as a molecular “brake” to synaptic strengthening (Anjum et al., 2024).

Despite this growing body of molecular and physiological data, it remains unclear how Rem2 directly influences learning and memory. In the present study, we used a conditional knockout approach to delete Rem2 specifically in CA1 neurons of the dorsal hippocampus and determine its role in spatial learning measured through a standard spatial object recognition **(SOR)** task. We then used an automated Western blot system (Li et al., 2020) to assess how Rem2 impacted the expression of all known AMPAR and NMDAR subunits in the CA1 of each mouse tested: GluA1, GluA2, GluA3, GluA4, GluN1, GluN2A, GluN2B, GluN2C, GluN2D, and GluN3A. This approach revealed an unexpected role of Rem2 in directly regulating GluN2D expression in CA1 interneurons of female but not male mice. Moreover, we found an indirect role of Rem2 in regulating the overall glutamate receptor composition of the CA1. These direct and indirect mechanisms likely give rise to the differences in spatial learning between male and female mice. To our knowledge, this is the first demonstration of a signaling molecule that confers sexual dimorphism to excitatory synapses in mammals. Understanding Rem2-dependent signaling could therefore offer important insights on how sex-dependent symptoms arise in neurodevelopmental and neurodegenerative disorders.

## 2 METHODS

### 2.1 Animals

Transgenic S129 mice carrying a floxed Rem2 allele (Moore et al., 2018) were bred in-house and maintained under standard laboratory conditions (12 h light/dark cycle, ad libitum access to food and water). Animals were group-housed by sex (maximum four per cage). All behavioral testing occurred during the light phase. Littermates were randomly assigned to experimental groups, which included conditional knockout (cKO) and sham controls. All procedures were conducted in accordance with the guidelines of the Institutional Animal Care and Use Committee (IACUC) at Temple University.

### 2.2 Stereotaxic injections

At postnatal day 28 (P28 ± 1), mice were anesthetized with isoflurane and placed in a stereotaxic frame. Following scalp incision and microdrill craniotomy, bilateral injections (0.2 μL per site) were made into the dorsal CA1 hippocampus using pulled glass pipettes. Sham mice were injected with pAAV.hSyn.eGFP (Addgene - #50465-AAV5), while cKO mice received pENN.AAV.hSyn.HI.eGFP-Cre.WPRE.SV40 (Addgene – #105540-AAV5). Coordinates relative to bregma were: ±1.4 mm ML, −1.95 mm AP, −1.15 mm DV. Meloxicam was administered subcutaneously immediately after surgery and again 48 hours later. Mice were allowed to recover for 14 days post-injection to ensure sufficient viral expression, Cre-mediated recombination, and Rem2 protein knockdown (Fig. 1A).

**Figure 1.**
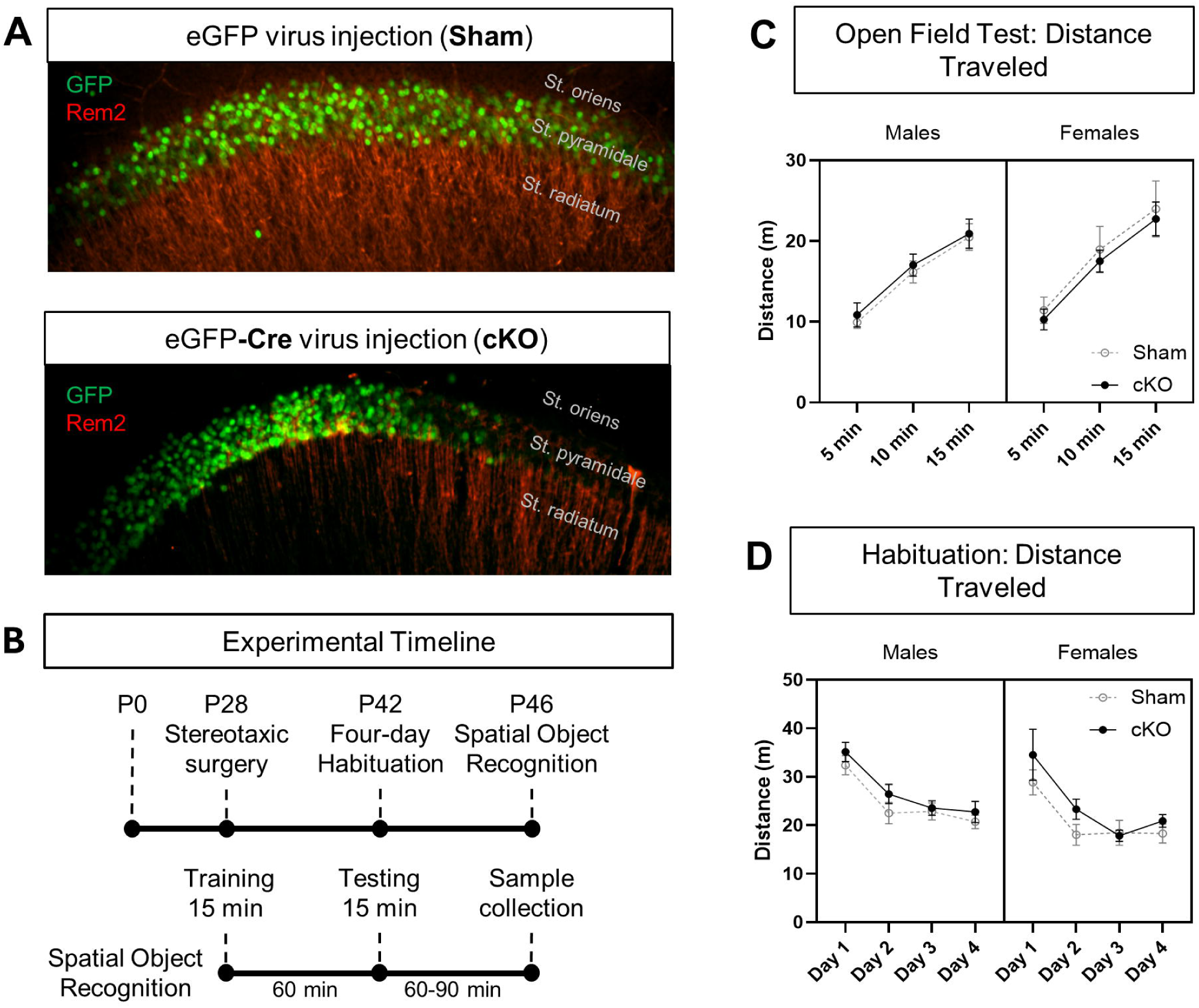
Experimental Design to Assess the Role of Rem2 in Spatial Memory. (A) Dorsal CA1 brain slices with AAV infected cells labeled with GFP and anti-Rem2 antibody. Rem2 knockout (Bottom) shows a significant decrease compared to Sham (Top). (B) Timeline of stereotaxic surgery, arena habituation, and overview of SOR. (C) Distance traveled in 5-minute intervals did not change upon Rem2 deletion in males or females. Mean ± SEM. (Males: Sham n = 11, cKO n = 12; Females: Sham n = 9, cKO n = 10) (D) Distance traveled over four days of habituation did not change based upon Rem2 deletion. Mean ± SEM. (Males: Sham n = 11, cKO n = 12; Females: Sham n = 9, cKO n = 10)

### 2.3 Experimental design

Following recovery, mice underwent a five-day behavioral protocol (Fig. 1B). Day 1 included the Open Field Test **(OFT)**, which also served as the initial habituation trial. Days 2–4 consisted of continued habituation to reduce novelty-induced behavioral variability. On Day 5, mice completed the Spatial Object Recognition **(SOR)** task. Brains were collected 60 minutes post-testing and bisected; one hemisphere was flash-frozen for biochemical analysis, and the other fixed in 4% paraformaldehyde (PFA) for immunohistochemistry.

### 2.4 Behavioral Tests

#### 2.4.1 Open Field Test (OFT) and Habituation

The open field arena measured 40 × 40 × 35 cm, with grey metal floors and black acrylic walls. A triangular white visual cue was affixed to the north wall for spatial reference. Arenas were uniformly illuminated at 25 lux using diffuse white light. Animals were individually placed into the center of the arena and allowed to explore freely for 15 minutes. The arena was digitally subdivided into a 5 × 5 grid (25 squares, each 80 × 80 mm). OFT data were analyzed in three 5-minute time bins to assess temporal dynamics of behavior. Habituation was conducted by repeating OFT for three additional days (Days 2–4) under identical conditions.

#### 2.4.2 Spatial Object Recognition (SOR)

SOR testing occurred on Day 5 in the same arena as the OFT. The assay consisted of three phases: (1) a 15-minute training session in which mice were exposed to two identical objects (2-inch plastic bottles) located in the northeast and northwest corners of the arena; (2) a 60-minute delay in the home cage; and (3) a 15-minute test session in which one object was relocated to either the southeast or southwest corner. Objects and arenas were cleaned with 70% ethanol between trials. Exploration was recorded and analyzed using ANY-maze software (Stoelting). Object investigation was defined as time spent facing the object within 25 mm, excluding climbing or rearing on the object. A discrimination index (DI) was calculated as: DI = (Time investigating moved object – Time investigating stationary object) / Total investigation time.

### 2.5 Tissue preparation

Mouse brains were harvested 60 minutes after behavioral testing. One hemisphere was flash-frozen for Western blots and the other was fixed for immunohistochemistry.

#### 2.5.1 Automated Western Blot System

Western blotting was performed using the Jess™ automated capillary electrophoresis system (ProteinSimple). Dorsal CA1 regions were microdissected (Hagihara et al., 2009) from the flash-frozen hemispheres. Briefly, CA1 tissues from each mouse were separately homogenized in RIPA buffer containing protease inhibitors, and protein concentrations were determined via BCA assay. Samples were prepared at a concentration of 0.05 mg/mL and run using a 5 μL loading volume. Reactions were performed using the RePlex total protein normalization method, with results expressed as protein abundance normalized to total protein. The following antibodies were used to assess all known AMPAR and NMDAR subunits in the mammalian brain: GluA1 (Cell Signaling 13185S), GluA2 (Abcam ab133477), GluA3 (Cell Signaling 4676S), GluA4 (Abcam ab1508), GluN1 (Thermofisher 32-0500), GluN2A (Cell Signaling 4205S), GluN2B (Cell Signaling 4207S) GluN2C (Novus NB300-107), GluN2D (Abcam ab314664), GluN3A (Abcam ab302534).

#### 2.5.2 Immunohistochemistry (IHC)

Hemispheres fixed in 4% paraformaldehyde were embedded in agarose and sectioned at 50 μm using a Compresstome VF-500-OZ (Precisionary Instruments). Free-floating sections underwent antigen retrieval in sodium citrate buffer (pH 8.0) at 80°C for 30 minutes, followed by blocking in 10% normal goat serum with 0.02% Triton X-100. Sections were incubated in primary antibodies (Rem2 C-11 Santa Cruz sc-514999 1:100) for 36 hours at 4°C, followed by secondary antibody incubation for 3 hours at room temperature. To reduce autofluorescence, sections were treated with 1:20 diluted TrueBlack® (Biotium) lipofuscin quencher in ethanol, then mounted and imaged at 20× magnification using an Olympus IX83 fluorescence microscope.

### 2.7 Data analysis

All behavioral data were recorded and quantified using ANY-maze (Stoelting). Western blot and behavioral data were analyzed in GraphPad Prism using two-way or three-way ANOVA, with Fisher’s LSD post hoc tests to assess group differences. Significance was defined as *P* ≤ 0.05.

### 2.8 Single-cell RNA sequencing data

Single-cell RNA-sequencing data were obtained from the Allen Institute mouse whole-brain 10x v3 dataset (Zeng-Aging-Mouse-10Xv3; metadata releases 20241130 and 20250131). Two expression matrices were used throughout: a pre-computed log2-normalized matrix for single-cell-level statistics, and the corresponding raw UMI count matrix for pseudobulk quantification. Both were read into R as SingleCellExperiment objects using the zellkonverter package. Donor sex categories were obtained from the accompanying cell metadata file.

#### 2.8.1 CA1 and Hippocampal Interneuron/Glia Cell-Type Selection

CA1 pyramidal neurons were defined by direct inclusion of the Allen whole-mouse-brain taxonomy subclass “016 CA1-ProS Glut” (cell_cluster_mapping_annotations.csv), independent of anatomical region restriction, as this subclass is already CA1-specific. Hippocampal interneuron and glial populations were drawn from broader region-level calls (Hippocampal Interneurons, Astrocytes, Microglia, Vascular, and OPC), with interneuron subtypes further resolved into short display codes from their Allen taxonomy subtype label (PV, PV-Chandelier, SST, VIP, Sncg, LAMP5, LAMP5-LHX6). This yielded twelve target cell-type categories: CA1 Pyramidal, PV, PV-Chandelier, SST, VIP, Sncg, LAMP5, LAMP5-LHX6, Astro, Micro, OPC, and Vascular, spanning both age cohorts (110,431 cells pooled across 29 donors. A young adult subset of this same population (donor_age_category = “adult”) with 39,195 cells across 9 donors (21,208 Male cells from 4 donors, 17,987 Female cells from 5 donors) was additionally carried forward for the sex-comparison analysis.

#### 2.8.2 Rem2 and Grin Gene Panel and Expression Quantification

Based on results from Western analyses, expression was quantified specifically for Rem2 and three NMDAR subunits: Grin1, Grin2c, and Grin2d. Genes were matched to the expression matrices by Ensembl gene IDs. Single-cell-level expression was taken directly from the pre-computed log2-normalized matrix. Pseudobulk expression was computed by summing raw UMI counts per donor within each cell type (donor × cell-type combinations with fewer than 10 cells were excluded) and normalizing to log2 (counts-per-million + 1) using each pool’s own total library size.

#### 2.8.3 Statistical Comparison Across Cell Types And Between Sexes

Expression differences across the twelve cell types were assessed per gene using a Kruskal-Wallis test (cell types with fewer than 10 cells were excluded), followed by Benjamini-Hochberg (BH) false discovery rate (FDR) correction across the 4-gene panel. Pairwise cell-type differences were further tested using two-sided Mann-Whitney U (Wilcoxon rank-sum) tests for each cell-type pair per gene, with rank-biserial correlation reported as effect size and log2 fold-change computed from mean expression (0.01 pseudocount); BH correction was applied within each gene’s set of pairwise comparisons.

A second analysis compared male vs female expression of the same four genes across the same twelve cell types. Two complementary tests were applied to each gene × cell-type combination. At single-cell resolution, a two-sided Mann-Whitney U test compared male vs female cells directly, with rank-biserial correlation as effect size and log2 fold-change from mean expression (0.01 pseudocount); this test served as a descriptive check rather than the primary inferential result, given pseudoreplication at the single-cell level. The primary donor-level test used pseudobulk log2(CPM + 1) values: a two-sided Wilcoxon rank-sum test and a Welch’s t-test were computed for cell types with at least two donors per sex, and DESeq2 was additionally applied to raw pseudobulk counts (design: ∼ donor_sex, female set as the reference level; genes retained if in the panel or if ≥10 counts in at least two donors). Because this panel contains only four genes, DESeq2’s default parametric or local-regression dispersion trend could not be reliably estimated (the local-regression fallback produced “out of vertex space” errors for some cell types); a single mean dispersion (fitType = “mean”) was used instead. Wald-test contrasts (Male vs Female) were extracted at α = 0.05, with BH FDR correction applied. Donor coverage after the ≥10-cell pseudobulk filter ranged from 2 to 4 donors per sex per cell type.

#### 2.8.4 Data Visualization

Mean log2 expression and row-wise z-scored expression across gene × cell-type combinations were visualized as heatmaps (ggplot2, diverging blue-white-red scale for z-scores: low #2166AC, mid white, high #B2182B, midpoint 0). Dot plots encoded mean expression (color) and detection rate, defined as the fraction of cells with expression > 0 (point size). Kruskal-Wallis significance across cell types was additionally plotted as a −log10(BH-adjusted p) bar chart per gene. Pairwise log2 fold-change matrices (cell type × cell type) were generated per gene, with asterisks marking BH-adjusted pairwise Wilcoxon *P* < 0.05. Pseudobulk expression was visualized as boxplots with per-donor jitter points, arranged in a panel grid specific to each dataset’s comparison axis (ggplot2/patchwork).

#### 2.8.5 Cell-Type UMAP Visualization

For both male and female datasets, an all-cohort UMAP spanning every annotated cell type in the dataset (not restricted to the CA1/interneuron/glia panel used above, or to any single cohort or age group) was generated from the same processed single-cell objects used elsewhere in this codebase and rendered in a shared arrow-labeled plotting style (ggrepel connector labels on a light-gray background). A PCA (top variable genes by variance, IRLBA) followed by UMAP was computed via the scatter package on all profiled cells (Seurat). These serve as orientation figures alongside the per-gene statistical results.

#### 2.8.6 Software and Reproducibility

All analyses were performed in R. Key packages included data.table, ggplot2, Matrix, patchwork, and DESeq2^2^ (Wald test, BH-adjusted p-values). The dataset was additionally accessed via zellkonverter and SingleCellExperiment^1^ for reading AnnData (.h5ad) matrices, with scater and BiocSingular used for the all-cohort UMAP embedding, reticulate/Python (anndata) used to load a pre-built expression subset for that step, and ggrepel used for UMAP label placement in both datasets. Each dataset’s analysis was executed as an independent R subprocess (Rscript) with full memory isolation between steps, orchestrated by a shared step-runner script that injects dataset-specific input/output paths and can skip steps whose expected output files already exist, allowing resumable execution.

## 3 RESULTS

### 3.1 CA1-specific Rem2 Deletion Enhances Spatial Learning in a Sex-dependent Manner

We generated Rem2 conditional knockout mice **(cKO)** and compared them to sham littermates (Fig. 1A; see Methods) to determine the role of Rem2 in CA1-dependent behaviors (Fig. 1B). We first assessed whether deleting Rem2 in CA1 neurons affected general activity and anxiety-related behaviors using the open field test **(OFT)**. Across sham and cKO groups, there were no significant differences in total distance traveled, time spent mobile, or time spent immobile (Fig. 1C, Suppl Fig. 1), indicating that knocking out Rem2 in CA1 neurons does not affect baseline locomotion or anxiety-related behavior.

Mice then underwent a four-day OFT habituation protocol (Fig. 1D) to minimize novelty-related variability before testing on the spatial object recognition **(SOR)** task which measures CA1-dependent spatial learning. We found that male shams significantly outperformed female shams in the SOR task (Fig. 2A, 2B). CA1-specific Rem2 deletion did not impact SOR performance in male mice (Discrimination Index, mean ± SEM: Male Sham = 0.16 ± 0.06, *n* = 10; Male cKO = 0.09 ± 0.06, *n* = 11; *P* = 0.31) likely due to a behavioral ceiling effect (Fig. 2A, 2B). In contrast, female mice with CA1-specific Rem2 deletion exhibited improved spatial learning compared to their sham littermates (Fig. 2A) as shown by their significantly higher discrimination index (Female Sham = −0.03 ± 0.03, *n* = 8; Female cKO = 0.14 ± 0.04, *n* = 10; *P* ≤ 0.05) (Fig. 2B). Effectively, Rem2 CA1 cKO females performed as well in the SOR task as male sham and male Rem2 CA1 cKO mice (Fig. 2B). We found no significant differences in locomotor activity or total object investigation time (Fig. 2C, D) indicating that the improved performance of female cKO mice was specific to spatial learning rather than changes in locomotion or anxiety-related behavior.

**Figure 2.**
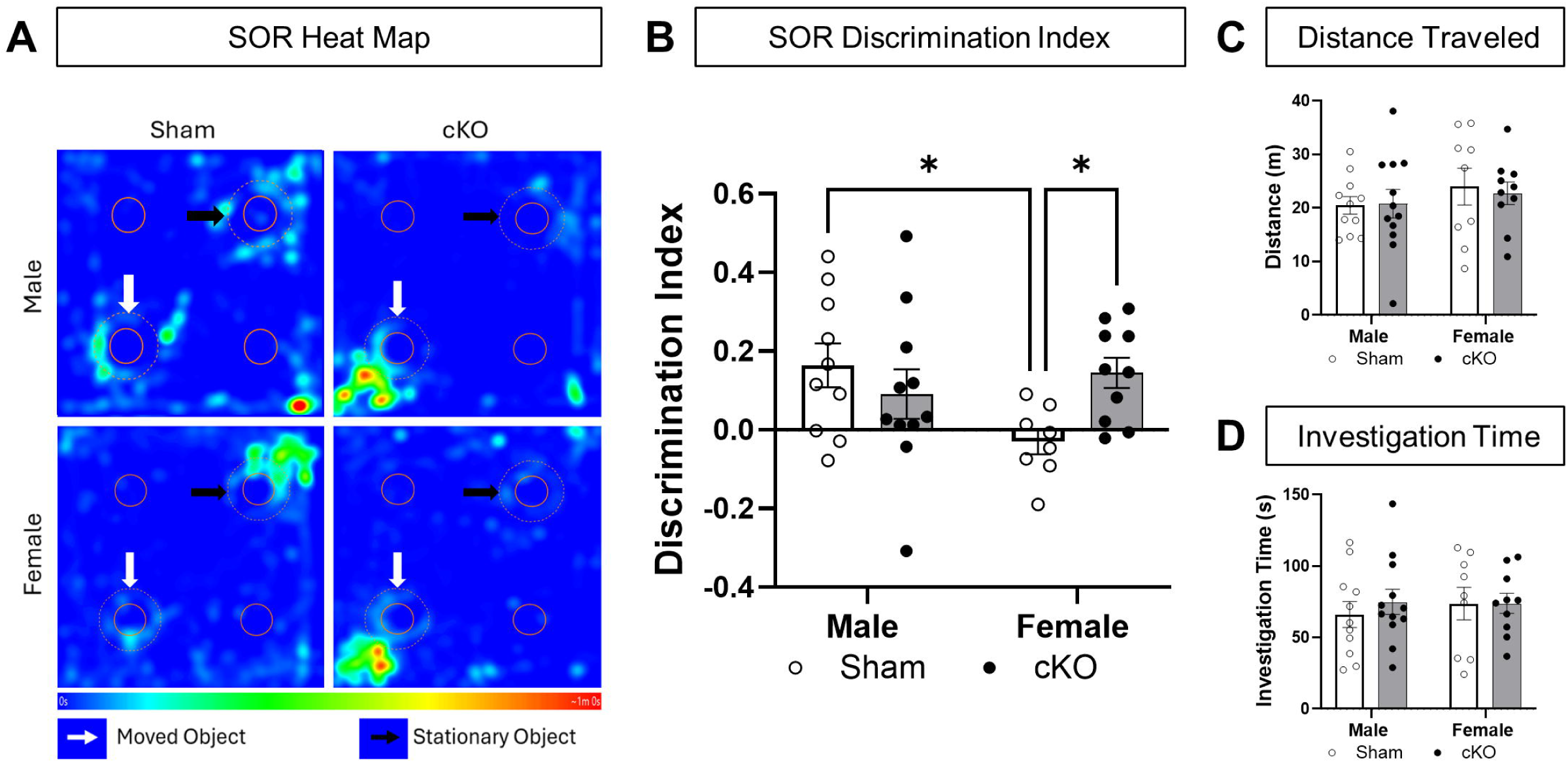
The Role of Rem2 in Spatial Learning. (A) Heat map indicating where mice spent time in the arena during the testing phase. (B) SOR discrimination index. Bar plots represent group data; circles represent individual animal data. Mean ± SEM. Two-way ANOVA; Interaction of Sex vs. Treatment F_1,35_ = 5.68, *P* < 0.05; Sex F_1,35_ = 1.82, *P* = 0.19; Treatment F_1,35_ = 5.68, *P* = 0.34. Male sham (n = 10) Female sham (n = 8) *P* < 0.05; Female sham (n = 8) Female cKO (n =10) *P* < 0.05; Fisher’s LSD post hoc test. (C) SOR distance traveled remained unchanged. Bar plots represent group data; circles represent individual animal data. Mean ± SEM. Two-way ANOVA; Interaction of Sex vs. Treatment F_1,35_ = 0.004, *P* = 0.95; Sex F_1,35_ = 1.12, *P* = 0.29; Treatment F_1,35_ = 0.004, *P* = 0.95. (D) SOR Investigation time remained unchanged. Bar plots represent group data; circles represent individual animal data. Mean ± SEM. Two-way ANOVA; Interaction of Sex vs. Treatment F_1,35_ = 0.02, *P* = 0.88; Sex F_1,35_ = 0.12, *P* = 0.73; Treatment F_1,35_ = 0.27, *P* = 0.61.

### 3.2 Rem2 Deletion Alters CA1 Glutamate Receptor Expression

To identify molecular correlates of our behavioral observations, we used an automated capillary-based Western blot system (Li et al., 2020; Okada et al., 2026) (Fig. 3A, 3B) to assess the expression of all known AMPA- and NMDA-mediated glutamate receptor (**AMPAR** and **NMDAR**) subunits (**GluA1-4**, **GluN1-3**) in every mouse that performed the SOR task. There were no AMPAR subunits that changed due to sex or as a result of CA1-specific Rem2 deletion in neurons (Fig. S2). In addition, neither GluN2A nor GlunN2B subunits, which are commonly incorporated in NMDARs to regulate synaptic plasticity (Sanz-Clemente et al., 2013; Ladagu et al., 2023), were altered in a sex- or Rem2-dependent manner (Fig. S2). However, we did find that both female sham and Rem2 CA1 cKO mice had significantly higher expression of the obligatory GluN1 subunit compared to their male counterparts (Female Sham, mean ± SEM = 2.21 ± 0.33, *n* = 8; Male Sham = 1.03 ± 0.11, *n* = 10; *P* ≤ 0.001; Female cKO = 2.10 ± 0.28, *n* = 10; Male cKO = 0.96 ± 0.08, *n* = 11; *P* ≤ 0.01) (Fig. 3C). Given that GluN1 expression is sex-dependent in both sham and Rem2 CA1 cKO mice, it alone is unlikely to account for our sex-dependent SOR results (Fig. 2A, B).

**Figure 3.**
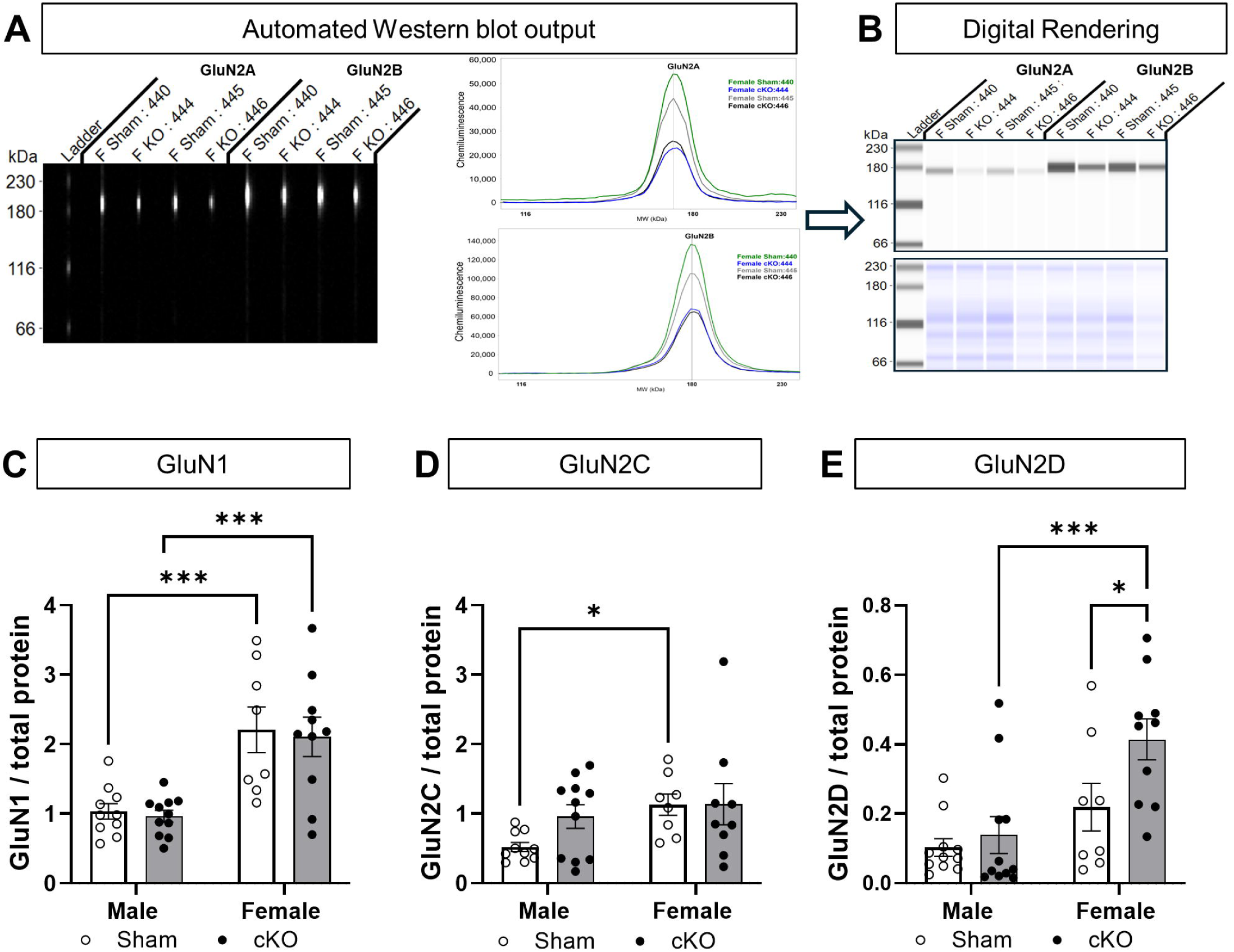
Assessment of AMPAR and NMDAR subunits in the CA1 subregion. (A) Representative capillary (left) and graphical (right) output of the Jess automated western blot machine. (B) Digital rendering shows traditional western blot view. (C) Bar plots represent group data for GluN1 expression; circles represent individual animal data. Mean ± SEM. Two-way ANOVA; Interaction of Sex vs. Treatment F_1,35_ = 0.01, *P* = 0.94; Sex F_1,35_ = 2.68, *P* < 0.0001; Treatment F_1,35_ = 2.68, *P* = 0.68. Male sham (n = 10) Female sham (n = 8) *P* < 0.001; Female sham (n = 8) Female cKO (n =10) *P* < 0.001; Fisher’s LSD post hoc test. (D) Bar plots represent group data for GluN2C expression; circles represent individual animal data. Mean ± SEM. Two-way ANOVA; Interaction of Sex vs. Treatment F_1,35_ = 1.34, *P* = 0.26; Sex F_1,35_ = 4.39, *P* < 0.05; Treatment F_1,35_ = 1.43, *P* = 0.24. Male sham (n = 10) Female sham (n = 8) *P* < 0.05; Fisher’s LSD post hoc test. (E) Bar plots represent group data for GluN2D expression; circles represent individual animal data. Mean ± SEM. Interaction of Sex vs. Treatment F_1,35_ = 5.68, *P* < 0.05; Sex F_1,35_ = 1.82, *P* = 0.19; Treatment F_1,35_ = 5.68, *P* = 0.34; two-way ANOVA. Male sham (n = 10) Female sham (n = 8) *P* < 0.05; Female sham (n = 8) Female cKO (n =10) *P* < 0.05; Fisher’s LSD hoc test.

Instead, the atypical GluN2C and GluN2D subunits appear to underlie our sex- and Rem2-dependent behavioral observations. Compared to NMDARs which commonly contain GluN2A and GluN2B subunits, NMDARs containing GluN2C and GluN2D are less sensitive to magnesium block and have slower deactivation time courses; however, they have lower single-channel conductance and are less permeable to calcium (Shelkar et al., 2019). Interestingly, we found a significant difference in GluN2C expression in male and female sham mice (Male Sham = 0.52 ± 0.07, *n* = 10; Female Sham = 1.13 ± 0.15, *n* = 8; *P* ≤ 0.05), but not in Rem2 CA1 cKO mice (Male cKO = 0.96 ± 0.17, *n* = 11; Female cKO = 1.14 ± 0.30, *n* = 9; *P* = 0.49) (Fig. 3D). In contrast, we found that deleting Rem2 in female mice significantly increased GluN2D expression compared to shams but did not affect GluN2D expression in males (Female Sham = 0.22 ± 0.07, *n* = 8; Female cKO = 0.41 ± 0.06, *n* = 10; *P* ≤ 0.05; Male Sham = 0.10 ± 0.03, *n* = 10; Male cKO = 0.14 ± 0.05, *n* = 11; *P* = 0.62) (Fig. 3E). Finally, we did not detect any significant changes in the seven remaining AMPAR and NMDAR subunits for either males or females (Fig. S2). These results indicate that Rem2-positive cells expressing GluN1, GluN2C, and/or GluN2D likely underlie our SOR results (Fig. 2).

### 3.3 Cell-specific action of Rem2

To determine if Rem2 acts on specific cell types to modulate sex-dependent spatial learning, we analyzed single-cell RNA sequencing (scRNA-seq) from the Allen Institute to determine what CA1 cells express *Rem2, Grin1, Grin2c,* and *Grin2d* (Fig. 4A; see Methods). We found no sex differences in the expression of these genes across the CA1 cell types we examined (Fig. 4B-E, Table S3).

**Figure 4.**
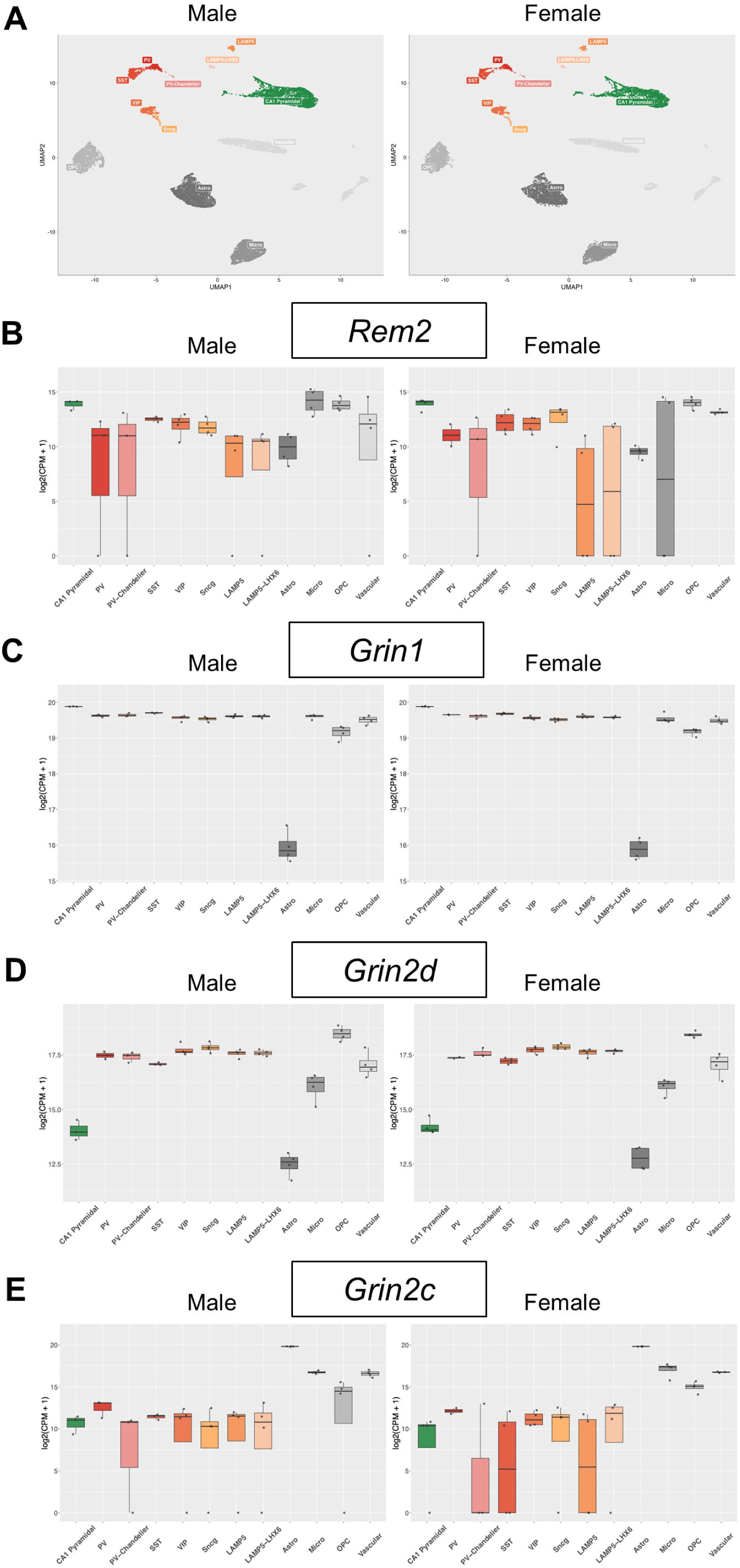
Cell-type expression patterns assessed from single cell RNAseq. (A) UMAPs of hippocampal CA1 cell-type data from Allen Institute. (B) CA1 cell-type expression levels of *Rem2* (C) CA1 cell-type expression levels of *Grin1* (D) CA1 cell-type expression levels of *Grin2d* (E) CA1 cell-type expression levels of *Grin2c*

We found that Rem2 was expressed in neuronal and non-neuronal cells including astrocytes, microglia, OPCs, and vascular cells (Fig. 4B). Among neurons, *Rem2* appears to be preferentially expressed in excitatory pyramidal cells but is also detectable in inhibitory neurons, most consistently in the SST, VIP, and Sncg subtypes (Fig. 4B).

We found that the *Grin1* gene was expressed similarly in all CA1 neuronal subtypes (Fig. 4C, Table S3). On the other hand, *Grin2d* was preferentially expressed in interneurons over CA1 pyramidal cells (Fig. 4D, Table S3) (Alsaad et al., 2018). These results suggest that sex-dependent effects of Rem2 on SOR (Fig. 2) are likely mediated by GluN2D-containing NMDARs in inhibitory interneurons (Fig. 4D). Finally, even though GluN2C is involved in sex differences in sham but not cKO mice (Fig. 3D), it is not likely targeted by Rem2 directly since *Grin2c* is preferentially expressed in astrocytes and non-neuronal cells (Fig. 4E) (Ravikrishnan et al., 2018; Alsaad et al., 2019).

### 3.4 Impact of Rem2 on Global Glutamate Receptor Composition of the CA1

Given that Rem2 is a member of the RGK family of non-canonical Ras-like GTPases, it is likely that its actions are not restricted to only regulating GluN2D expression in CA1 interneurons. Indeed, we found that deleting Rem2 changes the correlation between the expression of various AMPAR and NMDAR subunits—even those unrelated to GluN2D—in a sex-dependent manner (Fig. 5A, B). Upon Rem2 deletion, male mice lost multiple correlations (GluN2A-GluA2, GluN1-GluA3, GluN2A-GluN1, GluN2B-GluN1, GluN2B-GluN2A, GluN2C-GluN2A) while gaining new ones (GluN1-GluA1, GluN1-GluA2, GluN2C-GluA1, GluA3-GluA2, GluN2B-GluA4, GluN2D-GluN2C, GluN3A-GluN2B). Female mice also lost (GluA2-GluA1, GluA3-GluA1, GluN2C-GluN1) and gained (GluA4-GluA3, GluN2A-GluN1, GluN2B-GluN1) correlated glutamate pairs.

**Figure 5.**
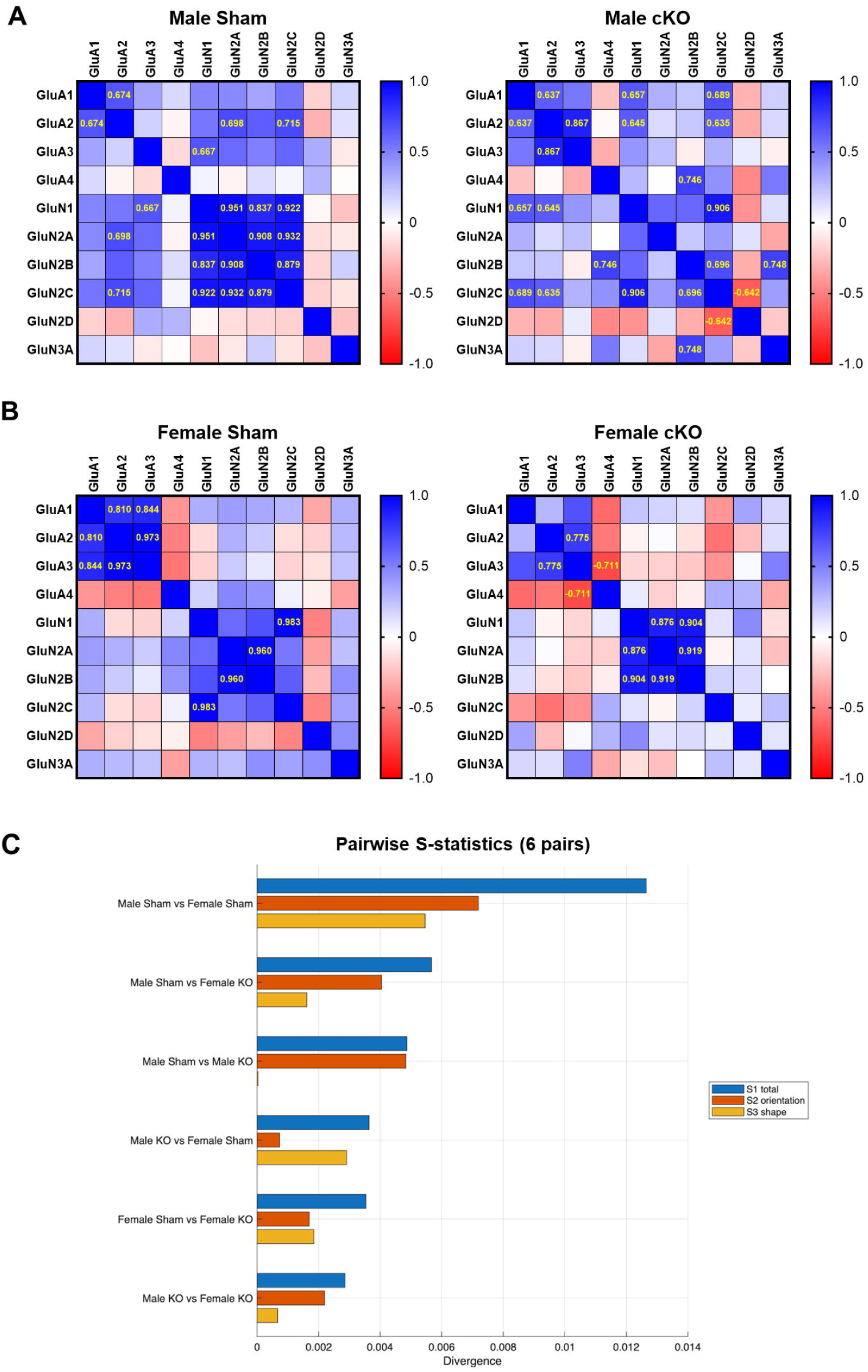
Effects on glutamate composition correlations. (A) Correlation matrix for male sham; Pearson’s coefficients given for significant (*P* = <0.05) pairs; n = 10) and male cKO; Pearson’s coefficients given for significant (*P* = <0.05) pairs; n = 11) (B) Correlation matrix for female sham; Pearson’s coefficients given for significant (*P* = <0.05) pairs; n = 7) and female cKO; Pearson’s coefficients given for significant (*P* = <0.05) pairs; n = 8) (C) Deletion of Rem2 reduced the divergence of Glutamate composition in males vs. females.

We used S-statistics to determine how these changes globally impact CA1 glutamate receptor composition among sham and Rem2 CA1 cKO, male and female mice (Fig. 5C; see Supplementary Methods). S-statistics compares how much variation in one dataset is captured by the principal component directions obtained from the other dataset. If the covariance structures are similar, each set of eigenvectors should explain similar amounts of variance in both datasets. The S-statistics are advantageous over standard principal component analysis (PCA) which analyzes one group at a time and cannot compare covariance structures across groups. The S-statistics are distribution-free, produce a continuous measure (*S*_1_), and *divide* it into orientation (*S*_2_) and shape (*S*_3_), making a targeted biological interpretation. However, a key limitation of S-statistics is that it does not have a formal theoretical framework of matrices’ properties (Garcia, 2012). Regardless, using S-statistics showed that the largest divergence (i.e. covariance) in global CA1 glutamate receptor composition occurred when comparing male sham versus female sham mice, which is similar to our SOR results (Fig. 2). Deleting Rem2 in CA1 neurons decreased this divergence by half such that all comparisons involving Rem2 CA1 cKO mice, male or female, were similar (Fig. 5C), much like our SOR results (Fig. 2). These findings suggest that Rem2 helps maintain sex differences in the overall glutamate receptor composition of the CA1 that could then dictate sexually dimorphic SOR behavior (Fig. 2).

## 4 Discussion

Identifying molecular regulators of synapse formation is critical for understanding and ultimately treating cognitive dysfunction due to neurodevelopmental and neurodegenerative disorders (Gadhave et al., 2024). In this study, we investigated how the small GTPase Rem2—a key regulator of excitatory synapse formation—impacts learning and memory. While prior studies have established the role of Rem2 in experience-dependent processes like ocular dominance and LTP, its direct role in cognitive behaviors is poorly understood.

Here we show that Rem2 deletion in the dorsal CA1 results in improved SOR performance in female mice only, indicating that Rem2 is a negative regulator of spatial learning (Fig. 2A, B). Rem2-specific deletion in CA1 neurons in male mice did not impact SOR, most likely due to a behavioral ceiling effect. These sex-dependent effects on SOR appears to be driven by the direct action of Rem2 on GluN2D-containing receptors in interneurons and its indirect action on the global glutamate receptor composition of the CA1. The latter mechanism could also reflect a compensatory response to Rem2 deletion. The fact that astrocytic GluN2C (Alsaad et al., 2019) was impacted by our neuron-targeted strategy (Fig. 1A) provides further evidence for potential homeostatic mechanisms in response to changes in CA1 synaptic activity as result of deleting Rem2.

GluN2C and GluN2D are atypical NMDAR subunits that—compared to GluN2A and GluN2B—are less sensitive to magnesium block, have slower deactivation time courses, lower single-channel conductance, and are less permeable to calcium. Some studies speculate that the presence of these receptors may have a neuroprotective effect on the network (Salimando et al., 2020; Camp et al., 2025; Sapkota et al., 2021). Our studies show an unexpected function of these receptors in maintaining sexually dimorphic behaviors.

A key limitation of our studies is that we did not observe any changes in SOR behaviors in males even though we found significant changes in their CA1 glutamate receptor composition as a result of Rem2 deletion. This is because, regardless of Rem2 expression levels, male mice could successfully perform our passive spatial task and thus there was little room for improvement (i.e. ceiling effect) unlike female mice. To thoroughly assess the behavioral impact of Rem2 therefore requires the use of other more active CA1-dependent tasks like contextual fear conditioning or active place avoidance.

## 5 Conclusion

This study presents the first demonstration of the effects of Rem2 on cognition identifying it as a negative regulator of CA1-dependent spatial learning. Our studies also show that the direct action of Rem2 on GluN2D expression as well as its indirect action on GluN2C could underlie its sex-dependent effects on spatial learning. Finally, our results expand on previous work implicating Rem2 as a molecular ‘brake’ on synaptic strengthening (Anjum et al., 2024) by presenting a model where deleting Rem2 could promote LTP by: 1) increasing GluN2D expression in interneurons of females causing increased synaptic inhibition between these cells (i.e. disinhibition of the CA1) and 2) reorganizing the overall glutamate receptor composition of the CA1 in both females and males. Interestingly, GluN2C and GluN2D expression have been independently shown to negatively modulate LTP (Dubois et al., 2015; Vestring et al., 2025). These findings underscore the need to elucidate the precise signaling pathways that enable Rem2, GluN2C, and GluN2D to regulate the identity of male and female synapses in the brain. Such studies are critical to our understanding of sex-dependent symptoms associated with neurodevelopmental and neurodegenerative disorders.

## Acknowledgements

This work was supported in part by a grant from the Charles E. Kaufman Foundation to V.L. (New Investigator Research Grant KA2023-136491).

---

**Figure S1.**
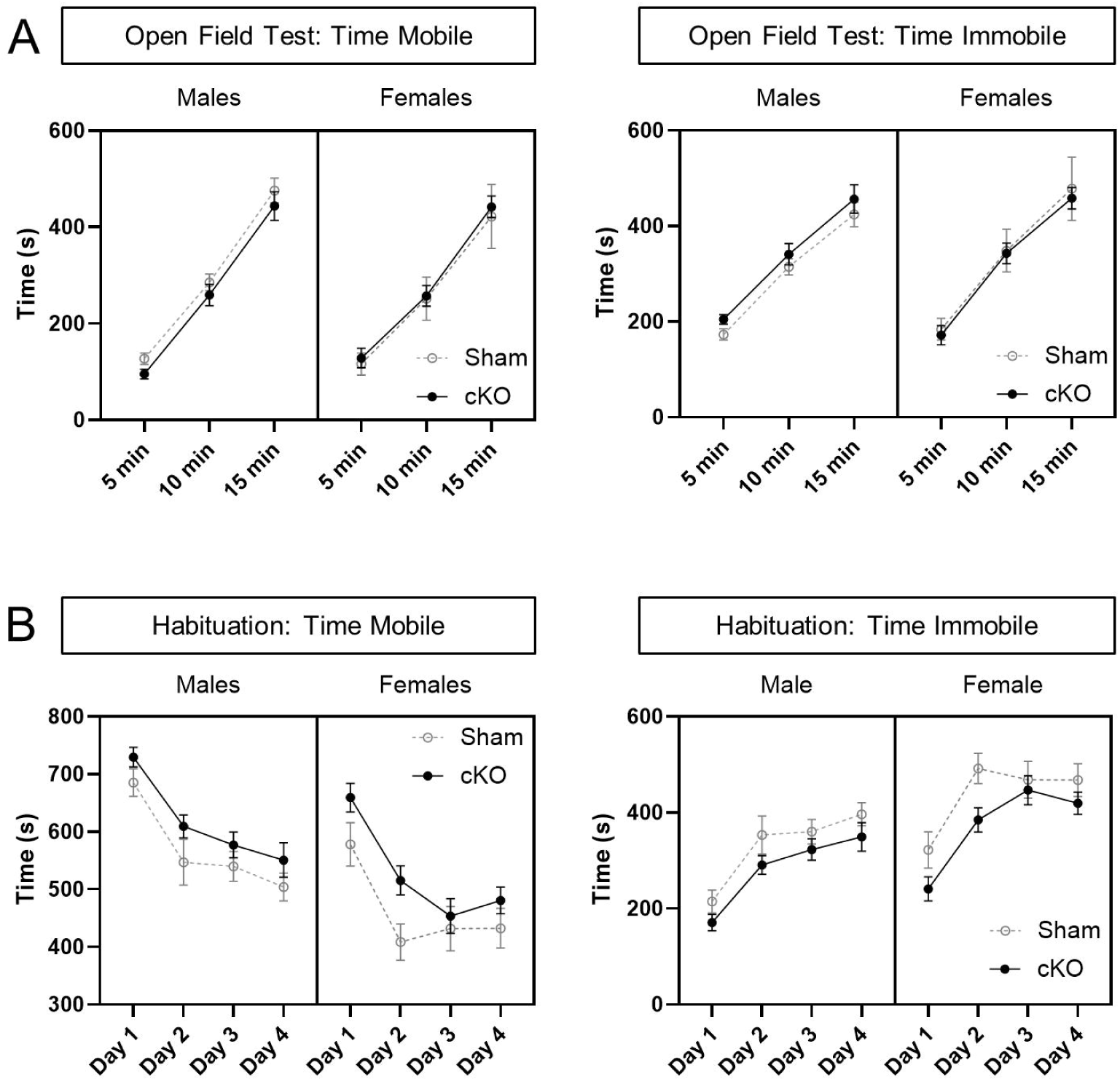
Mobility evaluations. (A) There were no changes in anxiolytic mobility over the duration of the OTF in either sex based on the presence of Rem2. (B) Both sexes exhibited reduced anxiolytic mobility during habituation. Day F_1.89,72.09_ = 34.67, *P* < 0.0001; sex F_1,38_ = 4.60, *P* < 0.05; treatment F_1,38_ = 3.38, *P* = 0.074; Mixed effects analysis.

**Figure S2.**
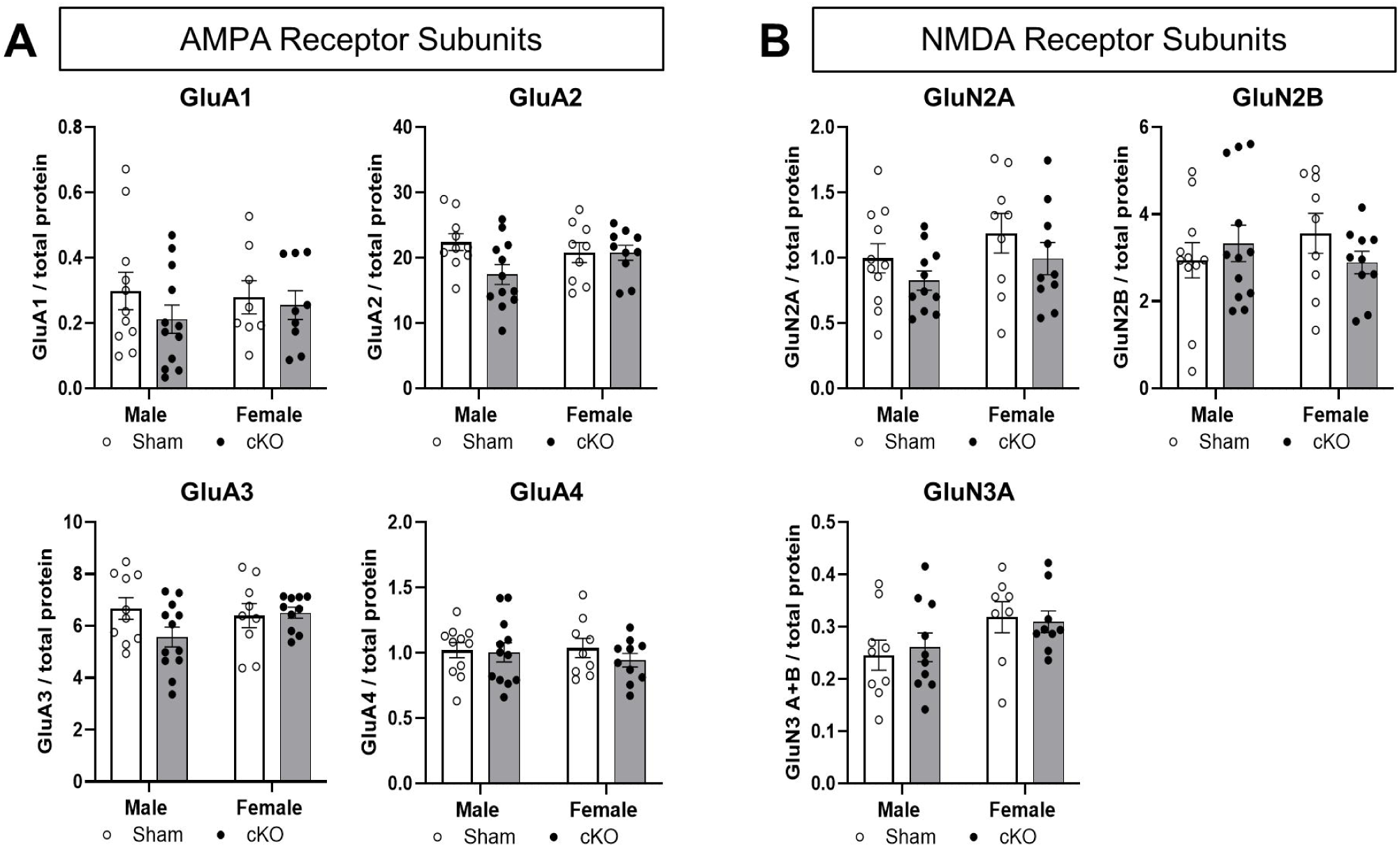
Glutamate receptors unaffected by Rem2 deletion. (A) No change in GluA1, GluA2, GluA3, or GluA4 expression due to the presence of Rem2. (B) No change in GluN2A, GluN2B, or GluN3A expression due to the presence of Rem2.

**Table S3.**
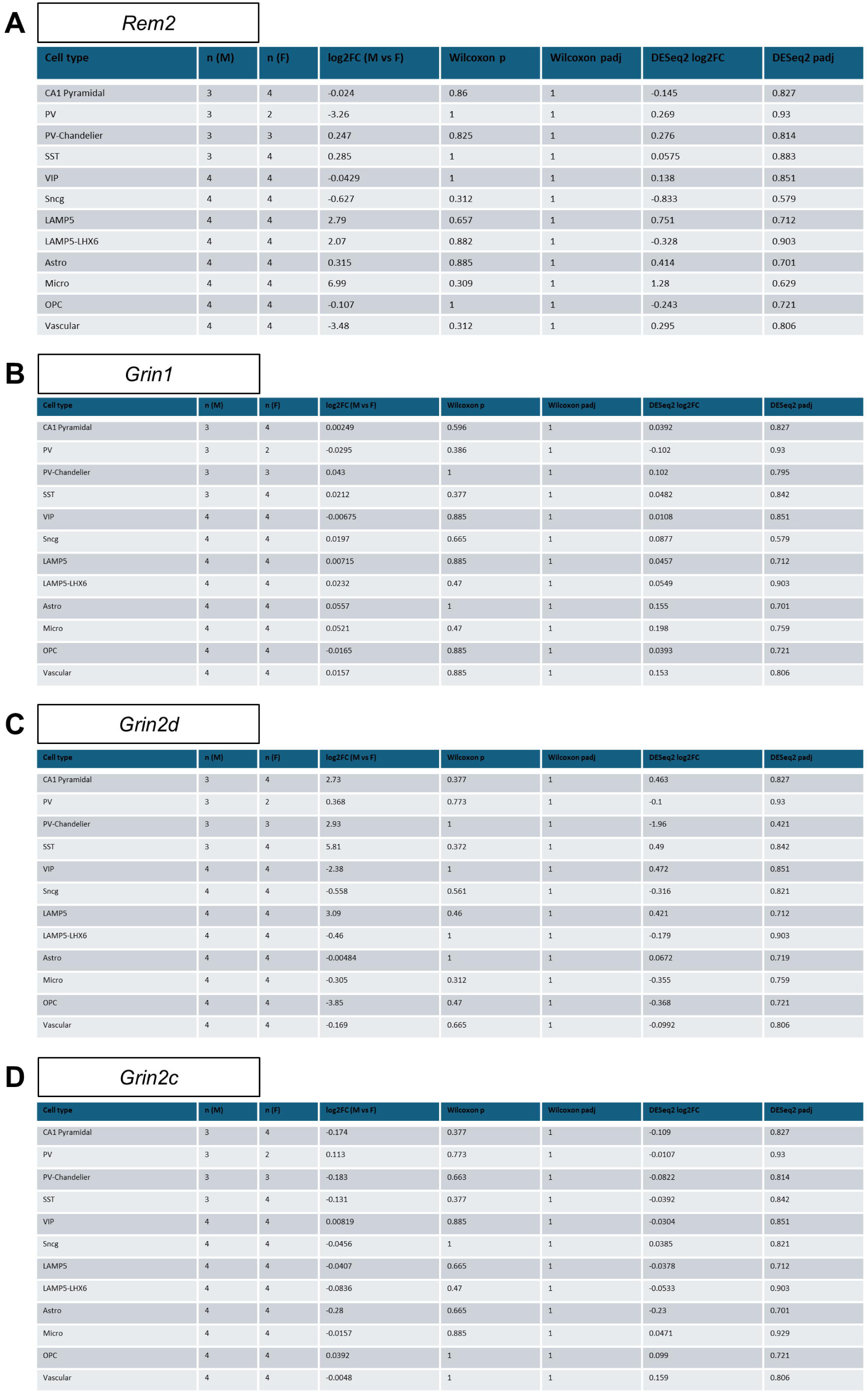
Quantification of single cell RNAseq cell-type expression data. (A) CA1 Cell-type expression data for *Rem2* (B) CA1 Cell-type expression data of *Grin1* (C) CA1 Cell-type expression data *Grin2d* (D) CA1 Cell-type expression data *Grin2c*

| Cell type | n (M) | n (F) | log2FC (M vs F) | Wilcoxon p | Wilcoxon padj | DESeq2 log2FC | DESeq2 padj |
| --- | --- | --- | --- | --- | --- | --- | --- |
| CA1 Pyramidal | 3 | 4 | -0.024 | 0.86 | 1 | -0.145 | 0.827 |
| PV | 3 | 2 | -3.26 | 1 | 1 | 0.269 | 0.93 |
| PV-Chandellier | 3 | 3 | 0.247 | 0.825 | 1 | 0.276 | 0.814 |
| SST | 3 | 4 | 0.285 | 1 | 1 | 0.0575 | 0.883 |
| VIP | 4 | 4 | -0.0429 | 1 | 1 | 0.138 | 0.851 |
| Sncg | 4 | 4 | -0.627 | 0.312 | 1 | -0.833 | 0.579 |
| LAMP5 | 4 | 4 | 2.79 | 0.657 | 1 | 0.751 | 0.712 |
| LAMP5-LHX6 | 4 | 4 | 2.07 | 0.882 | 1 | -0.328 | 0.903 |
| Astro | 4 | 4 | 0.315 | 0.885 | 1 | 0.414 | 0.701 |
| Micro | 4 | 4 | 6.99 | 0.309 | 1 | 1.28 | 0.629 |
| OPC | 4 | 4 | -0.107 | 1 | 1 | -0.243 | 0.721 |
| Vascular | 4 | 4 | -3.48 | 0.312 | 1 | 0.295 | 0.806 |

B *Grin1*
| Cell type | n (M) | n (F) | log2FC (M vs F) | Wilcoxon p | Wilcoxon padj | DESeq2 log2FC | DESeq2 padj |
| --- | --- | --- | --- | --- | --- | --- | --- |
| CA1 Pyramidal | 3 | 4 | 0.00249 | 0.596 | 1 | 0.0392 | 0.827 |
| PV | 3 | 2 | -0.0295 | 0.386 | 1 | -0.102 | 0.93 |
| PV-Chandellier | 3 | 3 | 0.043 | 1 | 1 | 0.102 | 0.795 |
| SST | 3 | 4 | 0.0212 | 0.377 | 1 | 0.0482 | 0.842 |
| VIP | 4 | 4 | -0.00675 | 0.885 | 1 | 0.0108 | 0.851 |
| Sncg | 4 | 4 | 0.0197 | 0.665 | 1 | 0.0877 | 0.579 |
| LAMP5 | 4 | 4 | 0.00715 | 0.885 | 1 | 0.0457 | 0.712 |
| LAMP5-LHX6 | 4 | 4 | 0.0232 | 0.47 | 1 | 0.0549 | 0.903 |
| Astro | 4 | 4 | 0.0557 | 1 | 1 | 0.155 | 0.701 |
| Micro | 4 | 4 | 0.0521 | 0.47 | 1 | 0.198 | 0.759 |
| OPC | 4 | 4 | -0.0165 | 0.885 | 1 | 0.0393 | 0.721 |
| Vascular | 4 | 4 | 0.0157 | 0.885 | 1 | 0.153 | 0.806 |

C *Grin2d*
| Cell type | n (M) | n (F) | log2FC (M vs F) | Wilcoxon p | Wilcoxon padj | DESeq2 log2FC | DESeq2 padj |
| --- | --- | --- | --- | --- | --- | --- | --- |
| CA1 Pyramidal | 3 | 4 | 2.73 | 0.377 | 1 | 0.463 | 0.827 |
| PV | 3 | 2 | 0.368 | 0.773 | 1 | -0.1 | 0.93 |
| PV-Chandellier | 3 | 3 | 2.93 | 1 | 1 | -1.96 | 0.421 |
| SST | 3 | 4 | 5.81 | 0.372 | 1 | 0.49 | 0.842 |
| VIP | 4 | 4 | -2.38 | 1 | 1 | 0.472 | 0.851 |
| Sncg | 4 | 4 | -0.558 | 0.561 | 1 | -0.316 | 0.821 |
| LAMP5 | 4 | 4 | 3.09 | 0.46 | 1 | 0.421 | 0.712 |
| LAMP5-LHX6 | 4 | 4 | -0.46 | 1 | 1 | -0.179 | 0.903 |
| Astro | 4 | 4 | -0.00484 | 1 | 1 | 0.0672 | 0.719 |
| Micro | 4 | 4 | -0.305 | 0.312 | 1 | -0.355 | 0.759 |
| OPC | 4 | 4 | -3.85 | 0.47 | 1 | -0.368 | 0.721 |
| Vascular | 4 | 4 | -0.169 | 0.665 | 1 | -0.0992 | 0.806 |

D *Grin2c*
| Cell type | n (M) | n (F) | log2FC (M vs F) | Wilcoxon p | Wilcoxon padj | DESeq2 log2FC | DESeq2 padj |
| --- | --- | --- | --- | --- | --- | --- | --- |
| CA1 Pyramidal | 3 | 4 | -0.174 | 0.377 | 1 | -0.109 | 0.827 |
| PV | 3 | 2 | 0.113 | 0.773 | 1 | -0.0107 | 0.93 |
| PV-Chandellier | 3 | 3 | -0.183 | 0.663 | 1 | -0.0822 | 0.814 |
| SST | 3 | 4 | -0.131 | 0.377 | 1 | -0.0392 | 0.842 |
| VIP | 4 | 4 | 0.00819 | 0.885 | 1 | -0.0304 | 0.851 |
| Sncg | 4 | 4 | -0.0456 | 1 | 1 | 0.0385 | 0.821 |
| LAMP5 | 4 | 4 | -0.0407 | 0.665 | 1 | -0.0378 | 0.712 |
| LAMP5-LHX6 | 4 | 4 | -0.0836 | 0.47 | 1 | -0.0533 | 0.903 |
| Astro | 4 | 4 | -0.28 | 0.665 | 1 | -0.23 | 0.701 |
| Micro | 4 | 4 | -0.0157 | 0.885 | 1 | 0.0471 | 0.929 |
| OPC | 4 | 4 | 0.0392 | 1 | 1 | 0.099 | 0.721 |
| Vascular | 4 | 4 | -0.0048 | 1 | 1 | 0.159 | 0.806 |

## References

Alsaad, H. A., DeKorver, N. W., Mao, Z., Dravid, S. M., Arikkath, J., & Monaghan, D. T. (2019). In the Telencephalon, GluN2C NMDA Receptor Subunit mRNA is Predominately Expressed in Glial Cells and GluN2D mRNA in Interneurons. Neurochem Res, 44(1), 61–77. doi:10.1007/s11064-018-2526-7

Anjum, R., Clarke, V. R. J., Nagasawa, Y., Murakoshi, H., & Paradis, S. (2024). Rem2 interacts with CaMKII at synapses and restricts long-term potentiation in hippocampus. bioRxiv. doi:10.1101/2024.03.11.584540

Arrant, A. E., Schramm-Sapyta, N. L., & Kuhn, C. M. (2013). Use of the light/dark test for anxiety in adult and adolescent male rats. Behav Brain Res, 256, 119–127. doi:10.1016/j.bbr.2013.05.035

Beguin, P., Ng, Y. J., Krause, C., Mahalakshmi, R. N., Ng, M. Y., & Hunziker, W. (2007). RGK small GTP-binding proteins interact with the nucleotide kinase domain of Ca2+-channel beta-subunits via an uncommon effector binding domain. J Biol Chem, 282(15), 11509–11520. doi:10.1074/jbc.M606423200

Boakye-Agyei, A. S., Rodrigues-Henry, D. D. M., Gonzalez, D. A., Sanders, E. M., & Dumas, T. C. (2026). Increased NMDA receptor GluN2A-type ionotropic signaling is sufficient to improve spatial memory in immature mice. Neurosci Lett, 878, 138595. doi:10.1016/j.neulet.2026.138595

Bourin, M., & Hascoet, M. (2003). The mouse light/dark box test. Eur J Pharmacol, 463(1-3), 55–65. doi:10.1016/s0014-2999(03)01274-3

Camp, C. R., Banke, T. G., Xing, H., Yu, K., Perszyk, R. E., Epplin, M. P., Akins, N.S., Zhang, J., Benke, T. A., Yuan, H., Liotta, D. C., Traynelis, S. F. (2025). Selective enhancement of the interneuron network and gamma-band power via GluN2C/GluN2D NMDA receptor potentiation. J Physiol, 603(14), 4027–4049. doi:10.1113/JP288343

Camp, C. R., & Yuan, H. (2020). GRIN2D/GluN2D NMDA receptor: Unique features and its contribution to pediatric developmental and epileptic encephalopathy. Eur J Paediatr Neurol, 24, 89–99. doi:10.1016/j.ejpn.2019.12.007

Chang, A., Liu, J. M., Nguyen, K., & Kumar, P. R. (2022). Expression optimization, purification, and biophysical characterization of a GluN2D-containing NMDA receptor. Protein Expr Purif, 198, 106129. doi:10.1016/j.pep.2022.106129

Chen, Q. Y., Li, X. H., Lu, J. S., Liu, Y., Lee, J. A., Chen, Y. X., Shi, W., Fan, K., Zhuo, M. (2021). NMDA GluN2C/2D receptors contribute to synaptic regulation and plasticity in the anterior cingulate cortex of adult mice. Mol Brain, 14(1), 60. doi:10.1186/s13041-021-00744-3

Dubois, C. J., Lachamp, P. M., Sun, L., Mishina, M., & Liu, S. J. (2016). Presynaptic GluN2D receptors detect glutamate spillover and regulate cerebellar GABA release. J Neurophysiol, 115(1), 271–285. doi:10.1152/jn.00687.2015

Eapen, A. V., Fernandez-Fernandez, D., Georgiou, J., Bortolotto, Z. A., Lightman, S., Jane, D. E., Volianskis, A., Collingridge, G. L. (2021). Multiple roles of GluN2D-containing NMDA receptors in short-term potentiation and long-term potentiation in mouse hippocampal slices. Neuropharmacology, 201, 108833. doi:10.1016/j.neuropharm.2021.108833

Finlin, B. S., Correll, R. N., Pang, C., Crump, S. M., Satin, J., & Andres, D. A. (2006). Analysis of the complex between Ca2+ channel beta-subunit and the Rem GTPase. J Biol Chem, 281(33), 23557–23566. doi:10.1074/jbc.M604867200

Finlin, B. S., Crump, S. M., Satin, J., & Andres, D. A. (2003). Regulation of voltage-gated calcium channel activity by the Rem and Rad GTPases. Proc Natl Acad Sci U S A, 100(24), 14469–14474. doi:10.1073/pnas.2437756100

Finlin, B. S., Mosley, A. L., Crump, S. M., Correll, R. N., Ozcan, S., Satin, J., & Andres, D. A. (2005). Regulation of L-type Ca2+ channel activity and insulin secretion by the Rem2 GTPase. J Biol Chem, 280(51), 41864–41871. doi:10.1074/jbc.M414261200

Finlin, B. S., Shao, H., Kadono-Okuda, K., Guo, N., & Andres, D. A. (2000). Rem2, a new member of the Rem/Rad/Gem/Kir family of Ras-related GTPases. Biochem J, 347 *Pt 1*(Pt 1), 223–231.

Fleischer, A. W., & Frick, K. M. (2023). New perspectives on sex differences in learning and memory. Trends Endocrinol Metab, 34(9), 526–538. doi:10.1016/j.tem.2023.06.003

Flynn, R., Chen, L., Hameed, S., Spafford, J. D., & Zamponi, G. W. (2008). Molecular determinants of Rem2 regulation of N-type calcium channels. Biochem Biophys Res Commun, 368(3), 827–831. doi:10.1016/j.bbrc.2008.02.020

Flynn, R., Labrie-Dion, E., Bernier, N., Colicos, M. A., De Koninck, P., & Zamponi, G. W. (2012). Activity-dependent subcellular cotrafficking of the small GTPase Rem2 and Ca2+/CaM-dependent protein kinase IIalpha. PLoS One, 7(7), e41185. doi:10.1371/journal.pone.0041185

Gadhave, D. G., Sugandhi, V. V., Jha, S. K., Nangare, S. N., Gupta, G., Singh, S. K., Dua, K., Cho, H., Hansbro, P. M., Paudel, K. R. (2024). Neurodegenerative disorders: Mechanisms of degeneration and therapeutic approaches with their clinical relevance. Ageing Res Rev, 99, 102357. doi:10.1016/j.arr.2024.102357

Ghiretti, A. E., Kenny, K., Marr, M. T., 2nd, & Paradis, S. (2013). CaMKII-dependent phosphorylation of the GTPase Rem2 is required to restrict dendritic complexity. J Neurosci, 33(15), 6504–6515. doi:10.1523/JNEUROSCI.3861-12.2013

Ghiretti, A. E., Moore, A. R., Brenner, R. G., Chen, L. F., West, A. E., Lau, N. C., Van Hooser, S. D., Paradis, S. (2014). Rem2 is an activity-dependent negative regulator of dendritic complexity in vivo. J Neurosci, 34(2), 392–407. doi:10.1523/JNEUROSCI.1328-13.2014

Ghiretti, A. E., & Paradis, S. (2011). The GTPase Rem2 regulates synapse development and dendritic morphology. Dev Neurobiol, 71(5), 374–389. doi:10.1002/dneu.20868

Ghiretti, A. E., & Paradis, S. (2014). Molecular mechanisms of activity-dependent changes in dendritic morphology: role of RGK proteins. Trends Neurosci, 37(7), 399–407. doi:10.1016/j.tins.2014.05.003

Hagihara, H., Toyama, K., Yamasaki, N., & Miyakawa, T. (2009). Dissection of hippocampal dentate gyrus from adult mouse. J Vis Exp(33). doi:10.3791/1543

Kenny, K., Royer, L., Moore, A. R., Chen, X., Marr, M. T., 2nd, & Paradis, S. (2017). Rem2 signaling affects neuronal structure and function in part by regulation of gene expression. Mol Cell Neurosci, 85, 190–201. doi:10.1016/j.mcn.2017.10.004

KK, S. N., & Dravid, S. M. (2026). GluN2D NMDA Receptors: Bridging Physiology and Pathology. Biol Psychiatry, 100(5), 493–507. doi:10.1016/j.biopsych.2026.02.012

Ladagu, A. D., Olopade, F. E., Adejare, A., & Olopade, J. O. (2023). GluN2A and GluN2B N-Methyl-D-Aspartate Receptor (NMDARs) Subunits: Their Roles and Therapeutic Antagonists in Neurological Diseases. Pharmaceuticals (Basel), 16(11). doi:10.3390/ph16111535

Li, Q., Jin, R., Zhang, S., Sun, X., & Wu, J. (2020). Group II metabotropic glutamate receptor agonist promotes retinal ganglion cell survival by reducing neuronal excitotoxicity in a rat chronic ocular hypertension model. Neuropharmacology, 170, 108016. doi:10.1016/j.neuropharm.2020.108016

Liput, D. J. (2018). Cre-Recombinase Dependent Germline Deletion of a Conditional Allele in the Rgs9cre Mouse Line. Front Neural Circuits, 12, 68. doi:10.3389/fncir.2018.00068

Liput, D. J., Lu, V. B., Davis, M. I., Puhl, H. L., & Ikeda, S. R. (2016). Rem2, a member of the RGK family of small GTPases, is enriched in nuclei of the basal ganglia. Sci Rep, 6, 25137. doi:10.1038/srep25137

Lloyd, J., & Litwa, K. (2026). Regulation of NMDA receptor interactions with the actin cytoskeleton in dendritic spine development. Front Cell Neurosci, 20, 1813072. doi:10.3389/fncel.2026.1813072

Lopes da Cunha, P., Villar, M. E., Ballarini, F., Tintorelli, R., & Ana Maria Viola, H. (2019). Spatial object recognition memory formation under acute stress. Hippocampus, 29(6), 491–499. doi:10.1002/hipo.23037

Lueptow, L. M. (2017). Novel Object Recognition Test for the Investigation of Learning and Memory in Mice. J Vis Exp(126). doi:10.3791/55718

Moore, A. R., Ghiretti, A. E., & Paradis, S. (2013). A loss-of-function analysis reveals that endogenous Rem2 promotes functional glutamatergic synapse formation and restricts dendritic complexity. PLoS One, 8(8), e74751. doi:10.1371/journal.pone.0074751

Moore, A. R., Richards, S. E., Kenny, K., Royer, L., Chan, U., Flavahan, K., Van Hooser, S. D., Paradis, S. (2018). Rem2 stabilizes intrinsic excitability and spontaneous firing in visual circuits. Elife, 7. doi:10.7554/eLife.33092

Okada, M., Okubo, R., Oka, T., & Motomura, E. (2026). Combined Inhibition of TRPM4/NMDA Receptor Complex and Extrasynaptic NMDA Receptors Is Candidate Therapeutic Target for Suppression of Epileptic Seizures and Improvement of Cognitive Impairments. Pharmacol Res Perspect, 14(3), e70256. doi:10.1002/prp2.70256

Olah, V. J., Witteveen, I. F., Banke, T. G., Leahy, A. M., Keable, R., Perszyk, R. E., Khayat, C. T., Nguyen, T. T., Bixler, B. J., Ghinger, F. G., Vullhorst, D., Yook, Y., Myers, S. J., Ullman, E. Z., Zhang, H., Traynelis, J. F., Diaz, E. S., Kim, S., Boraschi, S., Akins, N. S., Liu, K. H., Westhuyzen, A. V., Yang, Y., Camp, C. R., Buonanno, A., Hall, R. A., Zheng, J. Q., Liotta, D. C., Kennedy, M. J., Roche, K. W., Gourley, S. L., Yuan, H., Rowan, M. J. M., Traynelis, S. F. (2026). Bidirectional control of vesicular GABA release, LTP, and amotivation by GluN2D-selective allosteric modulators and ketamine. Cell Rep, 45(8), 117707. doi:10.1016/j.celrep.2026.117707

Paradis, S., Harrar, D. B., Lin, Y., Koon, A. C., Hauser, J. L., Griffith, E. C., Zhu, L., Brass, L. F., Chen, C., Greenberg, M. E. (2007). An RNAi-based approach identifies molecules required for glutamatergic and GABAergic synapse development. Neuron, 53(2), 217–232. doi:10.1016/j.neuron.2006.12.012

Ravikrishnan, A., Gandhi, P. J., Shelkar, G. P., Liu, J., Pavuluri, R., & Dravid, S. M. (2018). Region-specific Expression of NMDA Receptor GluN2C Subunit in Parvalbumin-Positive Neurons and Astrocytes: Analysis of GluN2C Expression using a Novel Reporter Model. Neuroscience, 380, 49–62. doi:10.1016/j.neuroscience.2018.03.011

Richards, S. E. V., Moore, A. R., Nam, A. Y., Saxena, S., Paradis, S., & Van Hooser, S. D. (2020). Experience-Dependent Development of Dendritic Arbors in Mouse Visual Cortex. J Neurosci, 40(34), 6536–6556. doi:10.1523/JNEUROSCI.2910-19.2020

Royer, L., Herzog, J. J., Kenny, K., Tzvetkova, B., Cochrane, J. C., Marr, M. T., 2nd, & Paradis, S. (2018). The Ras-like GTPase Rem2 is a potent inhibitor of calcium/calmodulin-dependent kinase II activity. J Biol Chem, 293(38), 14798–14811. doi:10.1074/jbc.RA118.003560

Salimando, G. J., Hyun, M., Boyt, K. M., & Winder, D. G. (2020). BNST GluN2D-Containing NMDA Receptors Influence Anxiety- and Depressive-like Behaviors and ModulateCell-Specific Excitatory/Inhibitory Synaptic Balance. J Neurosci, 40(20), 3949– 3968. doi:10.1523/JNEUROSCI.0270-20.2020

Sanz-Clemente, A., Nicoll, R. A., & Roche, K. W. (2013). Diversity in NMDA receptor composition: many regulators, many consequences. Neuroscientist, 19(1), 62–75. doi:10.1177/1073858411435129

Sapkota, K., Burnell, E. S., Irvine, M. W., Fang, G., Gawande, D. Y., Dravid, S. M., Jane, D. E., Monaghan, D. T. (2021). Pharmacological characterization of a novel negative allosteric modulator of NMDA receptors, UBP792. Neuropharmacology, 201, 108818. doi:10.1016/j.neuropharm.2021.108818

Schwartz, E. J., Rothman, J. S., Dugue, G. P., Diana, M., Rousseau, C., Silver, R. A., & Dieudonne, S. (2012). NMDA receptors with incomplete Mg(2)(+) block enable low-frequency transmission through the cerebellar cortex. J Neurosci, 32(20), 6878–6893. doi:10.1523/JNEUROSCI.5736-11.2012

Seibenhener, M. L., & Wooten, M. C. (2015). Use of the Open Field Maze to measure locomotor and anxiety-like behavior in mice. J Vis Exp(96), e52434. doi:10.3791/52434

Shelkar, G. P., Pavuluri, R., Gandhi, P. J., Ravikrishnan, A., Gawande, D. Y., Liu, J., Stairs, D. J., Ugale, R. R., Dravid, S. M. (2019). Differential effect of NMDA receptor GluN2C and GluN2D subunit ablation on behavior and channel blocker-induced schizophrenia phenotypes. Sci Rep, 9(1), 7572. doi:10.1038/s41598-019-43957-2

Vestring, S., Veleanu, M., Perez, M. C., Schuberth, L. E., Bronnec, M., Li, A., Wurz, L. M., Erdogdu, F., Stocker, J., Moos, J., Weigel, D., Theiss, A., Borger, L. M., Viota, S., Heynicke, F., Brandl, J., Hummel, F., Vivet, C., Jocher, D., Loewe, P., Barmann, S., Smoltczyk, L., Zimmermann, S., Prabhakaran, P., Lokaj, G., Sarrazin, D. H., Suarez, G., Bernhardt, J., du Vinage, C., Griessbach, E., Lais, J., Gensch, N., Wojtas, M., Knafo, S., Wendel, J., Warneke, J., Grohe, J. P., Gunter, S., Moumbock, A. F. A., Domschke, K., Serchov, T., Bischofberger, J., Normann, C. (2025). The NMDA receptor subunit GluN2D is a potential target for rapid antidepressant action. Nat Commun, 16(1), 10613. doi:10.1038/s41467-025-66774-w

Yang, T., & Colecraft, H. M. (2013). Regulation of voltage-dependent calcium channels by RGK proteins. Biochim Biophys Acta, 1828(7), 1644–1654. doi:10.1016/j.bbamem.2012.10.005

